# Putting computational models of immunity to the test - an invited challenge to predict *B. pertussis* vaccination outcomes

**DOI:** 10.1101/2024.09.04.611290

**Authors:** Pramod Shinde, Lisa Willemsen, Michael Anderson, Minori Aoki, Saonli Basu, Julie G Burel, Peng Cheng, Souradipto Ghosh Dastidar, Aidan Dunleavy, Tal Einav, Jamie Forschmiedt, Slim Fourati, Javier Garcia, William Gibson, Jason A Greenbaum, Leying Guan, Weikang Guan, Jeremy P Gygi, Brendan Ha, Joe Hou, Jason Hsiao, Yunda Huang, Rick Jansen, Bhargob Kakoty, Zhiyu Kang, James J Kobie, Mari Kojima, Anna Konstorum, Jiyeun Lee, Sloan A Lewis, Aixin Li, Eric F Lock, Jarjapu Mahita, Marcus Mendes, Hailong Meng, Aidan Neher, Somayeh Nili, Lars Rønn Olsen, Shelby Orfield, James A. Overton, Nidhi Pai, Cokie Parker, Brian Qian, Mikkel Rasmussen, Joaquin Reyna, Eve Richardson, Sandra Safo, Josey Sorenson, Aparna Srinivasan, Nicky Thrupp, Rashmi Tippalagama, Raphael Trevizani, Steffen Ventz, Jiuzhou Wang, Cheng-Chang Wu, Ferhat Ay, Barry Grant, Steven H Kleinstein, Bjoern Peters

## Abstract

Systems vaccinology studies have been used to build computational models that predict individual vaccine responses and identify the factors contributing to differences in outcome. Comparing such models is challenging due to variability in study designs. To address this, we established a community resource to compare models predicting *B. pertussis* booster responses and generate experimental data for the explicit purpose of model evaluation. We here describe our second computational prediction challenge using this resource, where we benchmarked 49 algorithms from 53 scientists. We found that the most successful models stood out in their handling of nonlinearities, reducing large feature sets to representative subsets, and advanced data preprocessing. In contrast, we found that models adopted from literature that were developed to predict vaccine antibody responses in other settings performed poorly, reinforcing the need for purpose-built models. Overall, this demonstrates the value of purpose-generated datasets for rigorous and open model evaluations to identify features that improve the reliability and applicability of computational models in vaccine response prediction.

## Introduction

Systems vaccinology aims to translate complex immunological data into actionable insights that can guide vaccination strategies. Achieving this requires integrating diverse datasets including genomic, proteomic, and transcriptomic data, to evaluate the systemic response to vaccination and build computational models of the vaccine-induced immune responses^1–3^. As a scientific community, we are advancing towards this goal by expanding cohort sizes, establishing meta-analyses involving a broad range of immune responses, and continuously integrating diverse datasets from single vaccines^4–6^ as well as multiple vaccines^7,8^ together. These efforts aim to capture the full complexity of the immune system and enhance our understanding of vaccine efficacy and safety across different populations ^4,8^.

A key challenge in this endeavor is to objectively test the generalizability and reproducibility of the findings generated by models developed in different studies. It is well known for genome-wide association studies^9^ that a given study can overemphasize dataset-specific results that do not replicate in other studies. The solution to this is to test previous findings in independent future studies. This can be challenging for systems vaccinology as there is significant variability between studies in terms of their design, specimen collection timing, and assays used to evaluate results. In addition, systems vaccinology studies are resource intensive, reducing the incentive for generating validation datasets. This means that most systems vaccinology-based models are generated based on datasets analyzed at the point of their publication, but they are not tested further on independent data.

To address this challenge, we initiated CMI-PB (Computational Models of Immunity to Pertussis Booster; https://www.cmi-pb.org). Our main goal is to test computational models that predict the outcome of booster vaccination which is performed through a series of data releases and associated community prediction contests. We have previously completed the first of three planned contests (**Table 1**) - a ‘dry-run’ involving CMI-PB consortium members forming teams using different models to answer the contest questions^10^. In the current study, we report our findings on the second ‘invited’ contest that included a select group of scientists from the broader community who have previously published in systems vaccinology. The datasets from a total of 96 subjects (**Table 1**) as part of the first challenge^10^ were made available as a training dataset to develop predictive models and we recruited a new cohort of 21 subjects which was available as an unseen testing dataset. We assessed over 49 computational models that applied various methodologies including classification-based techniques, such as naive Bayes and random forest, regression-based approaches like elastic net, and various other strategies encompassing multi-omics integration, gene signature analysis, and module scoring. We describe the details of these approaches, as well as general trends arising from the meta-analysis of all submissions. The full dataset, along with methods and scoring functions, are freely provided to the research community, and available to benchmark future algorithms in the field. The third public challenge will be open to community participation in August 2024.

**Table 1:** Overview of past and future CMI-PB prediction challenges. Our commitment involves conducting three annual challenges. The first challenge was completed in May 2022 with participation from the CMI-PB consortium. The second challenge concluded in January 2024 and featured the CMI-PB consortium along with a limited number of invited contestants from outside the consortium. We will involve members of the public in the third challenge. The second challenge included training data from the first challenge and newly generated challenge data. Similarly, we will use the training and challenge data from previous challenges as the training data for future challenges and generate new data for testing purposes.

|  | Prediction challenge title | Contestants | Subjects in dataset |  | Status |
| --- | --- | --- | --- | --- | --- |
|  |  |  | Train | Test |  |
| 1 | **First:** Internal dry run | CMI-PB consortium | 60 (28 aP + 32 wP) | 36 (19 aP + 17 wP) | Concluded in May 2022 |
| 2 | **Second:** Invited challenge | Invited contestants | 96 (47 aP + 49 wP) | 21 (11 aP + 10 wP) | Concluded in January 2024 |
| 3 | **Third:** Open Challenge | Public | 117 (58 aP + 59 wP) | 54 (27 aP + 27 wP) | Announced in August 2024 |

## Results

This results section covers two components: Sections 1-3 describe the experience of setting up and running the invited prediction contest. Sections 4-7 describe the specific models developed and discuss their performance on the prediction tasks.

### 1. Invitation of a select group of challenge participants

Our goal for this ‘invited challenge’ was to recruit external participants, but to keep the number at a manageable level of <50 teams to ensure we could provide individualized support. We consulted CMI-PB investigators and reviewed journal articles from Pubmed to identify a list of 50 scientists with prior experience in computational modeling, bioinformatics, or immunology and extended personal invitations to join the CMI-PB Challenge. Initially, 10 out of the 50 invited participants confirmed that they or their lab members would be interested while others mentioned conflicting schedules or time constraints as reasons for their inability to participate. Eventually, a total of 10 teams were formed, with three teams making up 5-6 PhD and masters students from the University of Minnesota each, one team of 3 researchers from different institutions, and the six teams remaining consisting of individual researchers for a total of 27 external participants. In addition to the invitations sent to external participants, we also invited participants from the labs of CMI-PB investigators who were not directly involved with the project, resulting in 8 participants, plus 1 team consisting of 5 master students from UCSD. Additionally, 5 members of the internal CMI-PB Consortium participated in the challenge. In total, we gathered 18 participating teams, for a total of 25 models submitted in the Challenge, which was a total of 49 people who participated in this invited challenge.

### 2. Summary of data sets and challenge tasks

#### Providing experimental data for training and testing prediction models

We generated data derived from more than 600 blood specimens collected from 117 subjects participating in a longitudinal study of *B. pertussis* booster vaccination. Blood specimens were collected on up to three days prior (day -30, -14, 0) and four days post-booster vaccination (day 1, 3, 7, and 14). The repeat pre-vaccination samples were intended to give a stable estimate of baseline and variability. For each specimen, we performed i) gene expression analysis (RNAseq) of bulk peripheral blood mononuclear cells (PBMC), ii) plasma cytokine concentration analysis, iii) cell frequency analysis of PBMC subsets, and iv) analysis of plasma antibodies against Tdap antigens (**Figure 1**; See Online Methods for a detailed description of the profiling data sets). The contestants were supplied with pre- and post-vaccination data as a training dataset to build their prediction models that consisted of two independent cohorts, the 2020 and 2021 cohorts, for a total of 96 subjects, which are discussed in detail in two previous publications^10,11^. For this challenge, we generated data from 21 new subjects. Baseline (pre-vaccination) challenge data was made available to contestants. The post-vaccine response challenge data was hidden from the contestants and used as ground truth for model evaluation.

**Figure 1.**
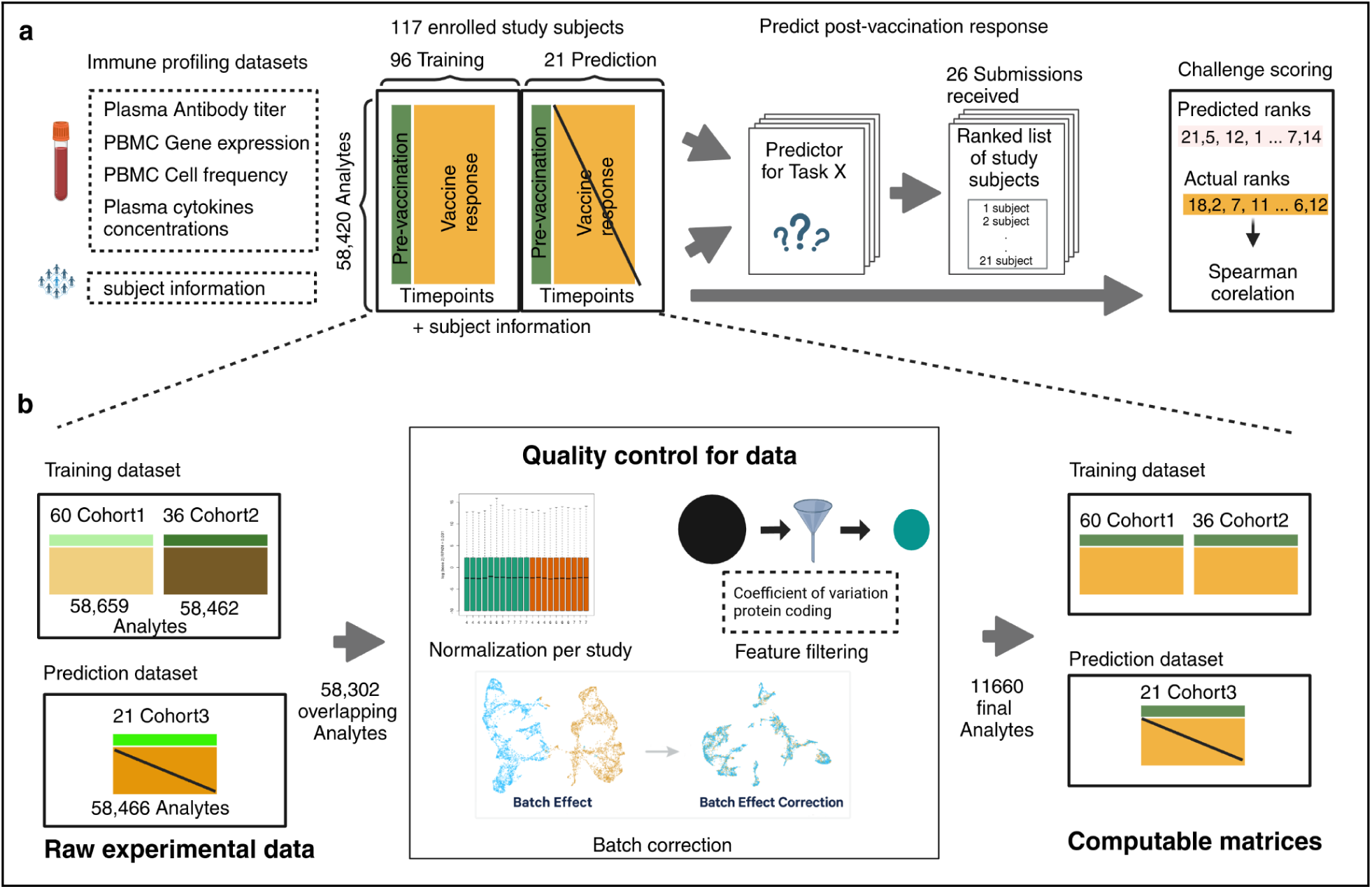
Generation of multi-omics datasets for 117 study participants. Contestants were provided with training datasets containing two cohorts (datasets 2020 and 2021), while the prediction dataset contained a newly generated cohort (dataset 2022). The training datasets contain pre-vaccination and post-vaccination immune response data, whereas the prediction dataset for 21 participants only contains pre-vaccination immune response data. Post-vaccination data was released after the challenge ended and used to evaluate submitted models.

#### Our data processing and harmonization approach

As the training dataset includes two multi-omics datasets from the 2020 and 2021 cohorts, which involved changes in the technicians performing the assay, and in the manufacturers setup of the assys, we are expecting batch effects that should be corrected before integrating them. While data processing and normalization methods are inherently user-specific, the CMI-PB team has developed a standardized data processing approach inspired by the methodology used in the internal CMI-PB challenge^10^. This involves 1) identifying common features, 2) baseline median normalization, and 3) batch-effect correction.

As a first step, we identified what features should be included in our analysis. Features are analytes measured in individual omics assays, such as cytokines in the Plasma cytokine concentrations assay. After the removal of features that were not found in all datasets, we were left with 58,302 overlapping features (**Figure 1A**). Many of these features had low information content, especially for the transcriptomic assay. To address this, for gene expression, we filtered zero variance and mitochondrial genes and removed lowly expressed genes (genes with transcript per million [TPM] <1 in at least 30% of specimens). Similarly, we filtered features with zero variance from cytokine concentrations, cell frequency, and antibody assays. This resulted in 11,660 features including 11,589 features from PBMC gene expression, 23 from PBMC cell frequency, 28 from Plasma cytokine concentrations, and 20 from Plasma antibody titers assays.

In the second step, we ran assay-specific data normalization. We performed baseline normalization on cell frequency, antibody titer, and cytokine concentration data. Specifically, we calculated the median using day zero time point data as a normalization factor per analyte and divided all values by this factor. We did not apply any normalization to the gene expression data. As a third step, we applied batch-effect correction on assay data within the training dataset to harmonize the data across 2020 and 2021 years. We employed the ComBat algorithm from the sva package, which adjusts for batch effects by modeling both batch and biological covariates^12,13^. After batch-effect correction, we validated the effectiveness of this step by examining the distribution of features across batches. We observed a significant reduction in cross-year batch-associated variability, confirming that the correction process was successful. This allowed us to move forward with a harmonized dataset for contestants for their analysis.

The challenge dataset underwent similar data processing and normalization to the training set to ensure consistency and comparability. This included using the median of pre-vaccination data to normalize cell frequency, antibody titer, and cytokine concentration data. We did not apply any normalization to the gene expression data. This processed data, along with raw data, was made available in TSV files and R data object formats, and the codebase used to transform from raw to processed was made available through GitHub.

#### Prediction tasks

We formulated six tasks asking contestants to predict a ranking of subjects from the highest response to the lowest response for each task based only on the pre-vaccination immune state data (**Table 2**). In task 1.1, contestants were asked to predict plasma IgG levels against the pertussis toxin (PT) on day 14 post-booster vaccination. Task 1.2 consisted of predicting the fold change of the plasma IgG levels against the pertussis toxin (PT) between day 14 post-booster vaccination and baseline. Tasks 2.1 and 2.2 required contestants to predict the overall frequency of monocytes among PBMCs on day 1 post-booster vaccination and the corresponding fold change, respectively. Similarly, in tasks 3.1 and 3.2 the CCL3 gene expression on day 3 post-booster vaccination and the corresponding fold change values compared to baseline needed to be predicted. This focus on 6 tasks that combine 3 targets with 2 readouts makes for a simpler setup compared to our previous competition.

**Table 2.**
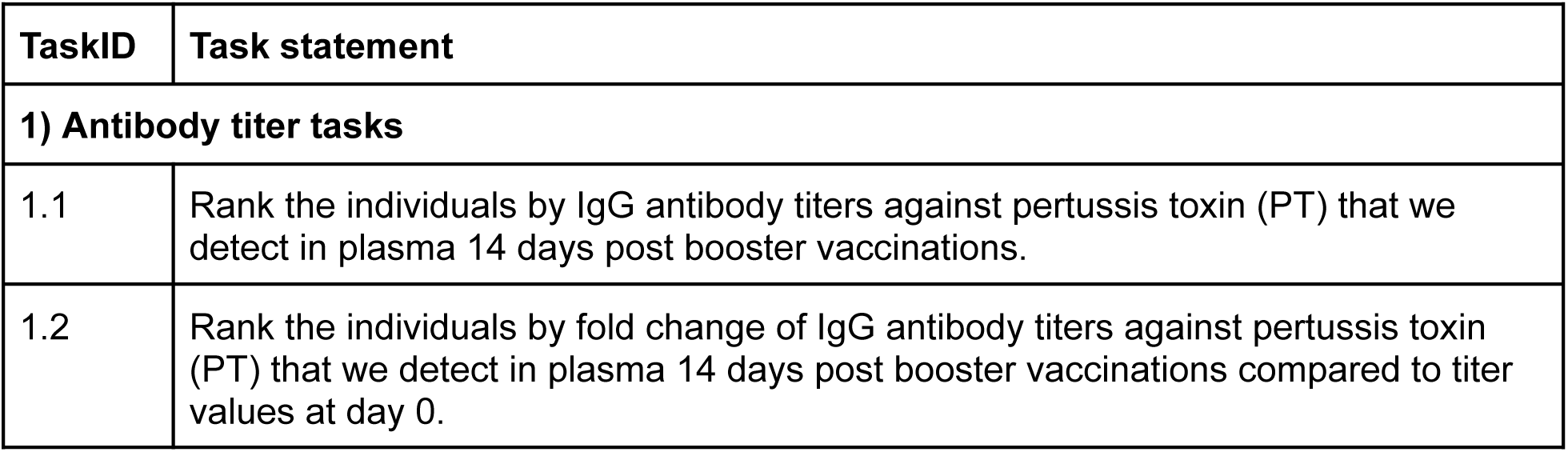
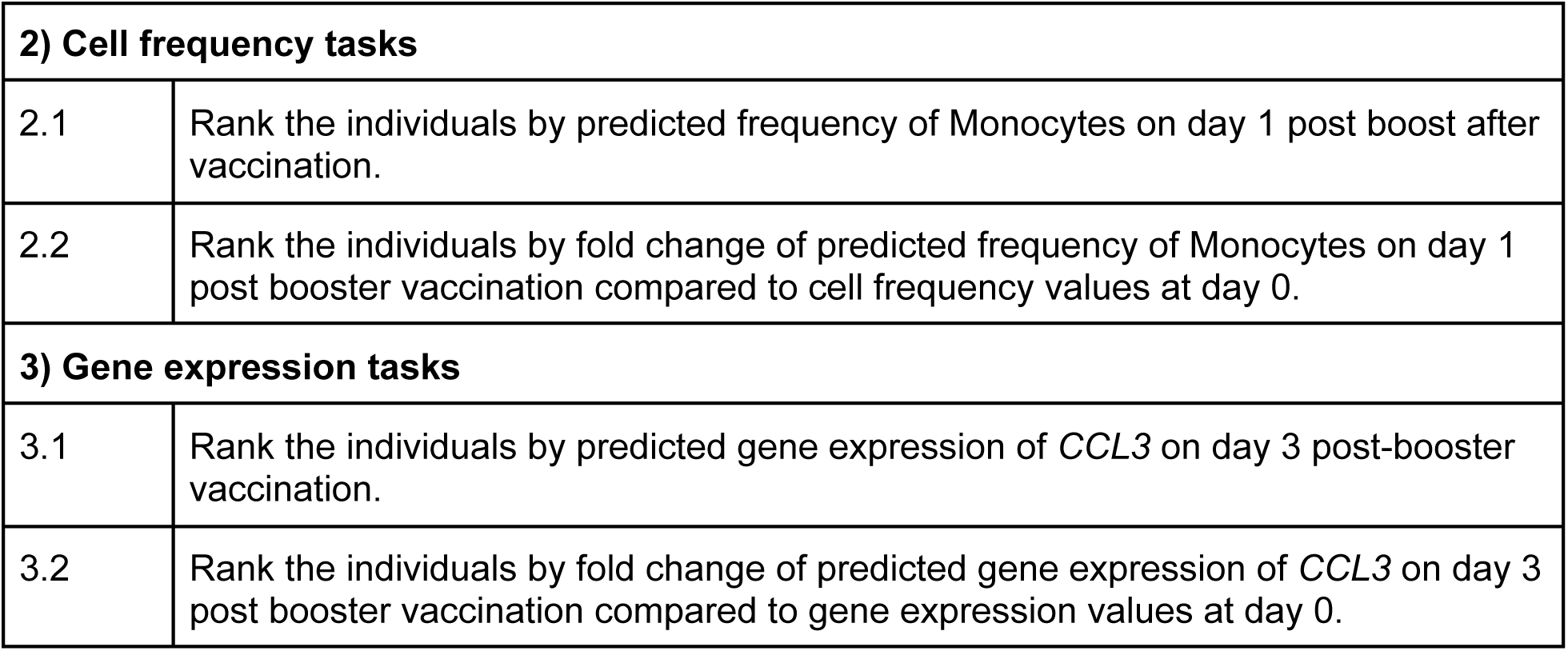
List of Prediction tasks. The tasks are grouped into three main types: antibody titer tasks, cell frequency tasks, and gene expression tasks. For each group, we asked to rank individual subjects based on either the absolute values of the biological readouts post-vaccination or the fold change compared to pre-vaccination measurement.

| TaskID | Task statement |
| --- | --- |
| **1) Antibody titer tasks** |  |
| 1.1 | Rank the individuals by IgG antibody titers against pertussis toxin (PT) that we detect in plasma 14 days post booster vaccinations. |
| 1.2 | Rank the individuals by fold change of IgG antibody titers against pertussis toxin (PT) that we detect in plasma 14 days post booster vaccinations compared to titer values at day 0. |

| 2) Cell frequency tasks |  |
| --- | --- |
| 2.1 | Rank the individuals by predicted frequency of Monocytes on day 1 post boost after vaccination. |
| 2.2 | Rank the individuals by fold change of predicted frequency of Monocytes on day 1 post booster vaccination compared to cell frequency values at day 0. |
| 3) Gene expression tasks |  |
| 3.1 | Rank the individuals by predicted gene expression of *CCL3* on day 3 post-booster vaccination. |
| 3.2 | Rank the individuals by fold change of predicted gene expression of *CCL3* on day 3 post booster vaccination compared to gene expression values at day 0. |

Each team could enter submissions for up to 3 different models and was allowed to update their submissions until the deadline. In total, We received 25 submissions for this invited challenge from 20 participating teams. In addition, we constructed 2 control models and incorporated 22 models previously identified from the literature, bringing the total number of models evaluated to 49. All teams provided detailed information about their computational methods and deposited their source code on GitHub, as listed in **Supplementary Note S1**.

### 3. Establishing control models and literature models

We established two simple control models that set a baseline of what more complex models should outperform. Model 1 is based on our finding that predicting vaccine responses solely based on the chronological age of the subject (the older, the worse) outperformed a lot of other models in predicting the antibody response to the Tdap vaccination^10^. Therefore, we implemented Control Model 1 simply by ranking subjects on their calendar age. Similarly, Control Model 2 captures that pre-vaccination levels of assay readouts are highly correlated with post-vaccination levels of the same readouts^14,15^. We implemented this for tasks 1.1 and 1.2, by using the baseline IgG antibody titer values against pertussis toxin as the predictor. For tasks 2.1 and 2.2, we used pre-vaccination monocyte frequencies, and for tasks 3.1 and 3.2, we used pre-vaccine levels of *CCL3* gene expression values. These control models are intended to set a baseline that more complex prediction models should exceed.

Additionally, we have implemented a set of 22 literature-derived models developed within the systems vaccinology field that aim to predict vaccination outcomes, as described in [38490204]. It is important to note that these models were repurposed for our prediction tasks and not evaluated in their original intended areas or studies. Instead, we evaluated these adapted models for their prediction performance on *B. pertussis* booster vaccination to determine the generalizability of these predictors. All of the literature models we identified were developed to predict antibody titers, so we only ran them on Task 1.

### 4. Contestants’ methods to predict vaccine response

We received a total of 25 submissions with the majority (19/25) of teams attempting all six tasks. Two teams completed five tasks, one team completed four tasks, two teams completed two tasks, and one team completed only one task. Contestants were asked to describe the methodologies they utilized, which included linear regression, nonlinear regression (regression trees), sparse linear regression, PLS (partial least-squares) or PC (principal component) regression, ensemble/model selection, and others (those methods not falling cleanly into the previous five categories). All methods are listed in **Table 3** with a short description that covers data pre- and postprocessing and expanded team summaries can be found in **Supplementary note S1**.

**Table 3.**
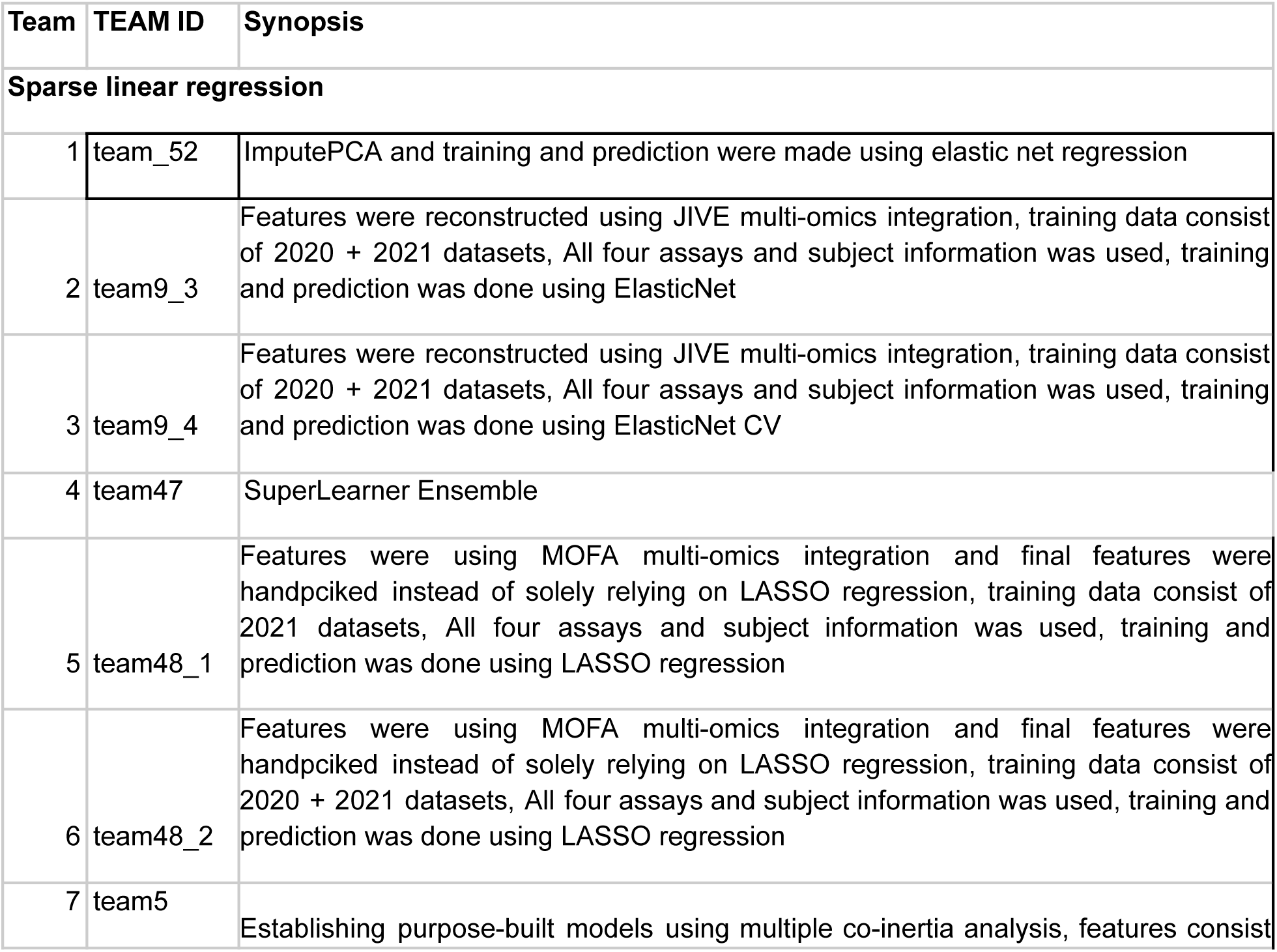
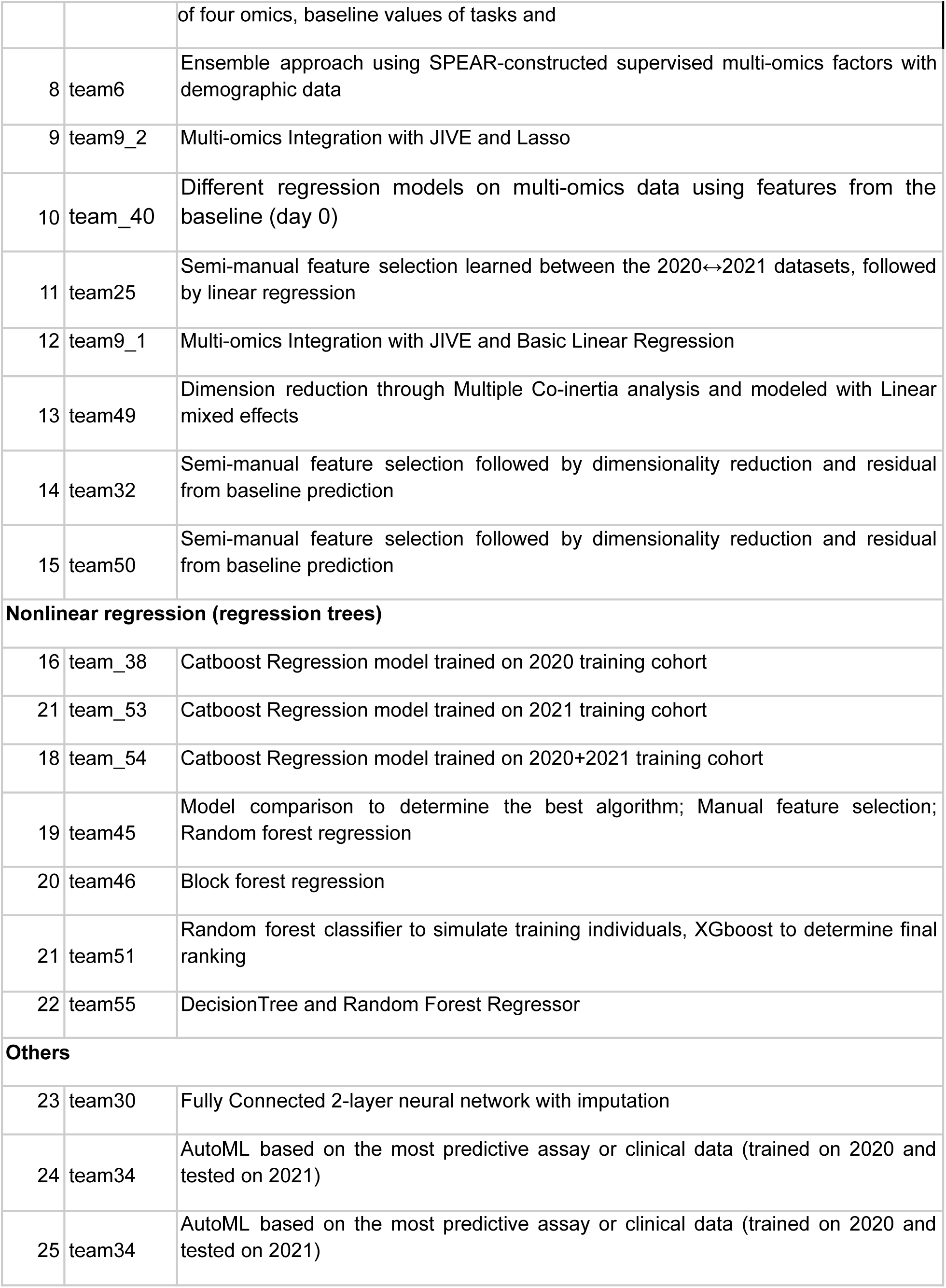
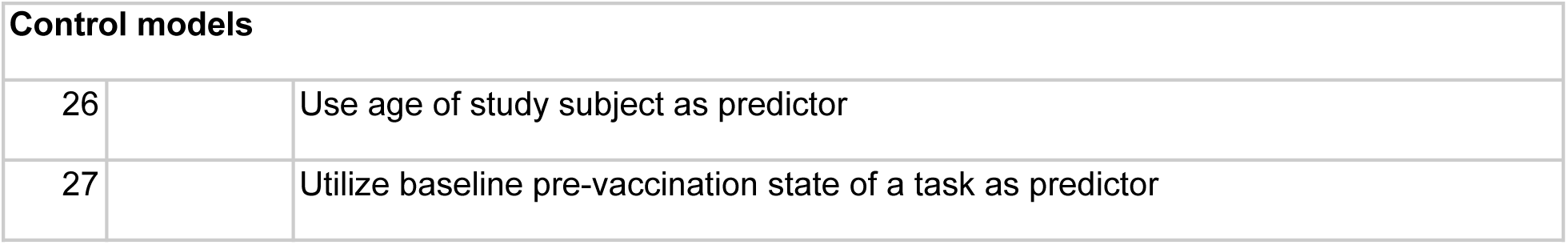
CMI-PB invited prediction challenge methods. The 25 team submissions were categorized according to their underlying methodology. Additional method characterizations can be found in **Supplemental Note 1**.

| Team | TEAM ID | Synopsis |
| --- | --- | --- |
| **Sparse linear regression** |  |  |
| 1 | team_52 | ImputePCA and training and prediction were made using elastic net regression |
| 2 | team9_3 | Features were reconstructed using JIVE multi-omics integration, training data consist of 2020 + 2021 datasets, All four assays and subject information was used, training and prediction was done using ElasticNet |
| 3 | team9_4 | Features were reconstructed using JIVE multi-omics integration, training data consist of 2020 + 2021 datasets, All four assays and subject information was used, training and prediction was done using ElasticNet CV |
| 4 | team47 | SuperLearner Ensemble |
| 5 | team48_1 | Features were using MOFA multi-omics integration and final features were handpciked instead of solely relying on LASSO regression, training data consist of 2021 datasets, All four assays and subject information was used, training and prediction was done using LASSO regression |
| 6 | team48_2 | Features were using MOFA multi-omics integration and final features were handpciked instead of solely relying on LASSO regression, training data consist of 2020 + 2021 datasets, All four assays and subject information was used, training and prediction was done using LASSO regression |
| 7 | team5 | Establishing purpose-built models using multiple co-inertia analysis, features consist |
|  |  | of four omics, baseline values of tasks and |
| 8 | team6 | Ensemble approach using SPEAR-constructed supervised multi-omics factors with demographic data |
| 9 | team9_2 | Multi-omics Integration with JIVE and Lasso |
| 10 | team_40 | Different regression models on multi-omics data using features from the baseline (day 0) |
| 11 | team25 | Semi-manual feature selection learned between the 2020↔2021 datasets, followed by linear regression |
| 12 | team9_1 | Multi-omics Integration with JIVE and Basic Linear Regression |
| 13 | team49 | Dimension reduction through Multiple Co-inertia analysis and modeled with Linear mixed effects |
| 14 | team32 | Semi-manual feature selection followed by dimensionality reduction and residual from baseline prediction |
| 15 | team50 | Semi-manual feature selection followed by dimensionality reduction and residual from baseline prediction |
| **Nonlinear regression (regression trees)** |  |  |
| 16 | team_38 | Catboost Regression model trained on 2020 training cohort |
| 21 | team_53 | Catboost Regression model trained on 2021 training cohort |
| 18 | team_54 | Catboost Regression model trained on 2020+2021 training cohort |
| 19 | team45 | Model comparison to determine the best algorithm; Manual feature selection; Random forest regression |
| 20 | team46 | Block forest regression |
| 21 | team51 | Random forest classifier to simulate training individuals, XGboost to determine final ranking |
| 22 | team55 | DecisionTree and Random Forest Regressor |
| **Others** |  |  |
| 23 | team30 | Fully Connected 2-layer neural network with imputation |
| 24 | team34 | AutoML based on the most predictive assay or clinical data (trained on 2020 and tested on 2021) |
| 25 | team34 | AutoML based on the most predictive assay or clinical data (trained on 2020 and tested on 2021) |

| Control models |  |  |
| --- | --- | --- |
| 26 |  | Use age of study subject as predictor |
| 27 |  | Utilize baseline pre-vaccination state of a task as predictor |

Most teams built their models using the provided preprocessed data. Some teams performed additional data processing required as a prerequisite for specific algorithms. These preprocessing techniques included data transformation and scaling (e.g., log10, square root), encoding for categorical features such as race and biological sex (e.g., label, one-hot), data imputation (e.g., PCA, Bayesian), and data normalization.

Preprocessing and feature selection are core components of building a predictor. In this challenge, features in the profiling data sets (P) far outnumber the total samples (N), increasing the risk of overfitting. To address this, teams often reduced the number of features modeled by correlating the features in the profiling dataset to the post-vaccination response data. A few teams also performed multi-omics integration and PC-based techniques to construct combined meta-features. Other preprocessing steps included principal component analysis, categorical regression, regularized regression (e.g., LASSO, ridge, or elastic nets), and mapping gene-level measurements to biological pathways or transcriptional modules.

Post-processing also differed in the specific models used for individual tasks. Most teams used summarizing or integrating one prediction model for all six tasks. In this approach, models were re-trained for specific tasks and evaluated separately to achieve better performance for each task. Other teams built entirely separate models for each task. Additionally, teams employed various cross-validation approaches, including leave-one-out, k-fold, 5-fold, and cross-cohort (testing on the 2020 cohort and evaluating on the 2021 cohort, and vice versa). Detailed descriptions of the team methods can be found in **Table 1**.

### 5. Evaluating Task performance on vaccine response predictions

We first evaluated the prediction performance of the control models and models from the literature. As specified in the contest description, Spearman’s Rank Correlation Coefficients were utilized as a metric for the evaluation of the submitted models for each task. For Control Model 1 which was solely based on the age of subjects, we found no significant relationship for any of the six tasks (**Figure 2A**). In contrast, we observed a significant positive correlation for Control Model 2 between the ranking of post-vaccination responses and their respective baselines for all three tasks: Monocytes on day 1, *CCL3* on day 3, and IgG-PT on day 14 (**Figure 2A**). This suggests that overall, the booster vaccination does not disrupt the pre-existing ranking of subjects in these readouts. In contrast, a strong negative correlation was noted between the fold change of IgG-PT at day 14 and its baseline. This translates to subjects with low pre-vaccination antibody titers showing the largest fold-change increase in titers post vaccination. Notably, this is not observed for the other two readouts (*CCL3* gene levels and Monocyte frequency), suggesting that it is not just a result of ‘regression to the mean’. Rather, individuals with very low antibody titers seem to benefit the most from a booster vaccination.

**Figure 2.**
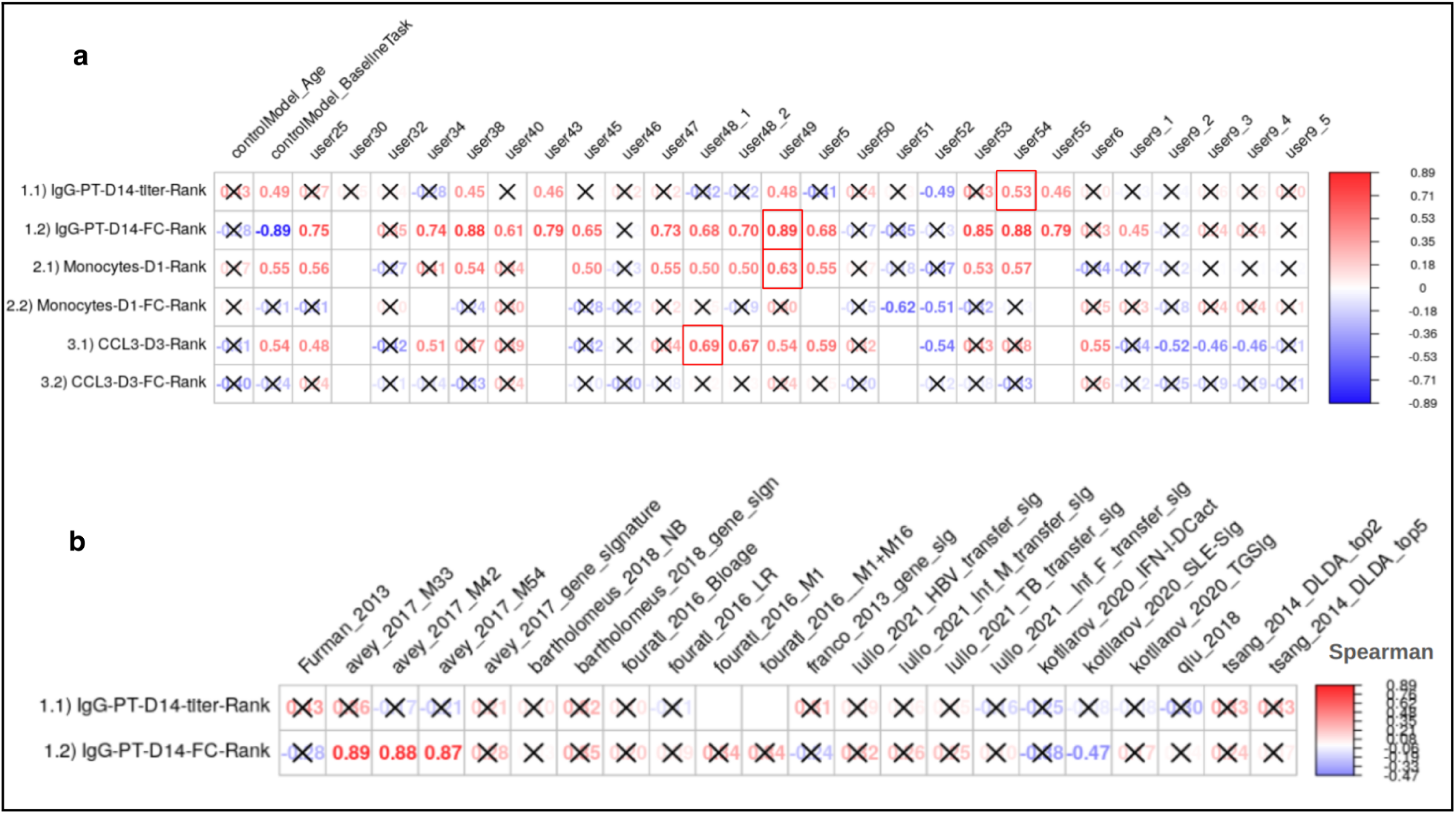
Evaluation of the prediction models submitted for the invited CMI-PB challenge. a) control models and models submitted by contestants b) models from systems vaccinology literature. Model evaluation was performed using Spearman’s rank correlation coefficient between predicted ranks by a contestant and actual rank for each of (1.1 and 1.2) antibody titers, (2.1 and 2.2) immune cell frequencies, and (3.1 and 3.2) transcriptomics tasks. The number denotes Spearman rank correlation coefficient, while crosses represent any correlations that are not significant using p ≥ 0.05.

Of the 22 literature models tested, only four provided a significant Spearman correlation coefficient, and all of those were for task 1.2 (antibody fold-change). None of the literature models outperformed the ‘baseline’ Control Model 2 (**Figure 2B**). Overall this suggests that the Control Models we implemented provided a good baseline that needs to be exceeded by new models to prove their value.

In terms of contestant-submitted predictions, among the 25 submissions received, 20 demonstrated at least one significant correlation coefficient. These models were considered important, and their performances are discussed subsequently (**Section 6**). In the top 20 models, prevalent techniques for selecting predictor genes included univariate feature ranking, meta-gene construction through multi-omics integration, and literature-based gene selection. The common prediction models employed were random forest and regularized regression methods (LASSO and ridge regression), with the latter being notably used by the top-ranked Team 49 in this sub-challenge.

Contestant-submitted predictions were aggregated by teams, where the score of each team was calculated using a point system to rank all submissions and identify the overall winner of the challenge. We awarded 3 points to the submission ranked highest in a particular task and 1 point if the contestant attempted the task. The team with the highest points was awarded as the winner of the challenge. The final scores revealed that the winning team is from the University of Minnesota (Team 49), achieving superior predictions in tasks 1.2 (r = 0.7, p-value = 0.001) and 2.1 (r = 0.81, p-value = 0.0031) (**Figure 2**). Two teams from the LJI (Teams 54 and 38) ranked second overall. A team from the National Institutes of Health (Team 51) ranked third overall and achieved the top rank for task 2.2 (see **Figure 2A** for details). Team 54 ranked top for task 1.1, and Team 38 ranked top for task 3.1. As no submissions showed a significant correlation coefficient for task 3.2, there was no team declared as winner for that task.

### 6. Top-performing methods include distinct approaches: multi-omics integration, categorical regression, and subject-based training

The top-performing team from the University of Minnesota developed a machine learning method that integrated multi-omics profiling data sets and knowledge-enhanced data representations into a nonlinear, probabilistic regression model to learn and predict vaccine response tasks (**Figure 3** and source code provided as Supplementary Software). Starting with raw experimental data, the workflow involved initial data imputation and batch effect correction that considered different time points separately to help maintain the temporal integrity of the data^16^. Feature selection was then performed using various statistical techniques, including LASSO, Ridge, PCA, PLS, and Multiple Co-Inertia Analysis (MCIA). MCIA was then chosen as the best-performing method, which integrates different data modalities to produce a reduced set of key multi-omics features^17^. These features were then utilized in a linear mixed effect model, structured to differentiate between fixed and random effects. The model was trained on a subset of the data, with validation through 5-fold cross-validation, and then tested to evaluate its predictive performance.

**Figure 3.**
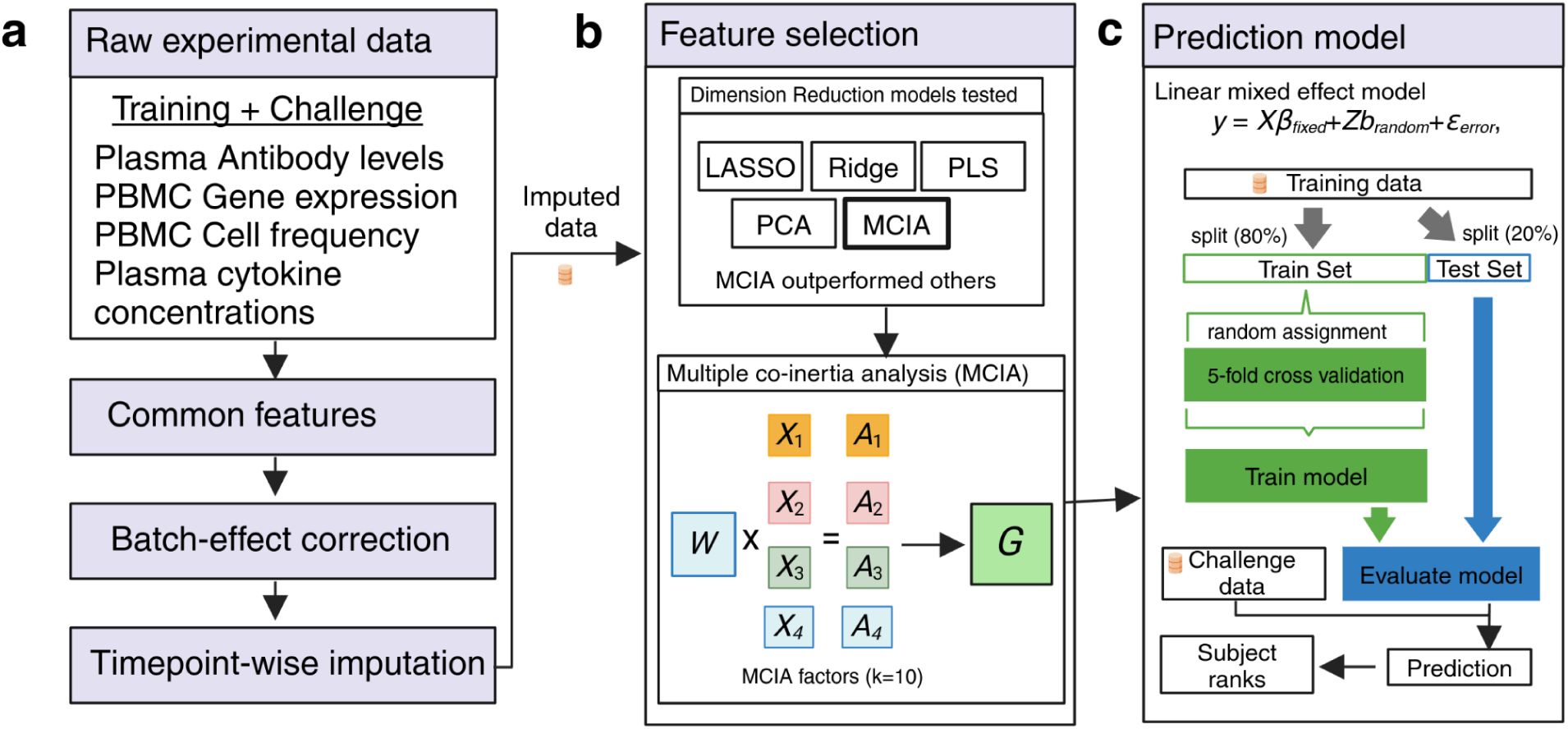
The method implemented by the winning team. Schematic overview of the data processing, feature selection, and prediction modeling workflow. (a) The workflow begins with raw experimental data, including training and challenge datasets from plasma antibody levels, PBMC gene expression, PBMC cell frequency, and plasma cytokine concentration assays. The common features across these datasets are identified, followed by batch-effect correction and timepoint-wise imputation. (b) Feature selection was performed using various dimension reduction techniques, including LASSO, Ridge, PLS, PCA, and Multiple Co-inertia Analysis (MCIA). MCIA outperformed the other models and was selected for further analysis. MCIA integrates different data types (e.g., X1, X2, X3, X4) and their associated weights (A1, A2, A3, A4) to produce MCIA factors (G) that represent the combined data structure. (c) These MCIA factors were then used in a Linear Mixed Effects (LME) model to predict the outcome. The model was trained on 80% of the data (train set) using 5-fold cross-validation and evaluated on the remaining 20% (test set). The trained model was then applied to the challenge baseline data to generate predictions, which were used to rank subjects according to their predicted outcomes.

There were two second-best-performing teams. The team led by Dr. Thrupp from LJI utilized multi-omics integration with Multi-Omics Factor Analysis (MOFA) which is also a factor analysis model that provides a general framework for the integration of multi-omic data sets in an unsupervised fashion^18^. Initially, processed data from the 2020 and 2021 training cohorts, which included all four assays, were used to construct 10 MOFA factors. Subsequently, LASSO was employed to identify the best-performing feature^19^. The model was trained on a subset of this data, validated through 5-fold cross-validation, and then tested to assess its predictive performance. The team led by Dr. Jarjapu from LJI utilized the Catboost Regression model, which was trained on the 2020 and 2021 training cohorts^20^. Feature selection was conducted manually, selecting features that exhibited consistent Spearman correlation coefficients when the model was trained separately on the 2020 and 2021 datasets. This approach ensured that only the most reliable and stable features were used for the final model, aiming to enhance the robustness and accuracy of the predictive outcomes.

The third-ranked team led by Dr. Gibson from the NIH adopted a distinctive strategy by employing a Random Forest classifier to simulate training individual subjects and XGBoost to determine the final rankings^21,22^. They attempted four of the six tasks, specifically excluding the gene expression tasks. This team utilized processed data from three assays: cell frequency, cytokine concentrations, and antibody titers. For tasks 1.1 and 1.2, they implemented data imputation using median antibody titers per feature. To validate their model, they applied K-fold validation, ensuring the robustness and reliability of their predictive model through systematic resampling and evaluation.

## Discussion

In this study, we evaluated multi-omics data from Tdap booster immunizations to predict vaccine outcomes. We focused on *Bordetella Pertussis (BP)* because of its continued public health importance and the ability to compare different vaccination regimes. BP causes whooping cough, a highly contagious respiratory infection that most severely affects infants^23^. The introduction of whole-cell pertussis (wP) vaccines in ∼1950 massively reduced the incidence of infections. Due to observed reactogenicity side effects, the wP vaccines were replaced with acellular pertussis (aP) vaccines in 1996. Following this, pertussis incidence has been rising in the last two decades, likely due to waning immunity post aP vaccination^24–28^. Studies, including our own^11,29,30^, have shown long-lasting effects and differences in T cell responses in adults originally vaccinated with aP versus wP vaccines, despite subsequent aP booster vaccination, but it remains unclear how these differences are maintained over time^31,32^. To address these questions, our near-term goal is to determine how an individual responds to pertussis antigen re-encounter by characterizing the resulting cascade of events (i.e., recall memory response) and relating it to the pre-vaccination immune state.

This ’invited’ challenge differed from our first ‘dry run’ challenge by including teams from labs other than the organizers. Insights gained from all 49 submitted methods and their relative performance provide a valuable resource for future algorithm development (**Table S2**). We observed that the top-performing methods employed distinct and innovative approaches to the challenge. These included strategies such as multi-omics data integration, which leverages the combined information from multiple omics to enhance predictive power; categorical regression, which effectively handles discrete outcome variables; and subject-based training, where models were tailored to individual-specific characteristics to improve accuracy in predicting vaccine responses. The diversity of these successful methodologies highlights the complex and multifaceted nature of TDap booster vaccination response prediction and emphasizes the importance of adopting various approaches to tackle this challenge effectively.

Furthermore, the presented results-based models showed significant Spearman correlation coefficients. Contestants employed diverse methods that included different composite features through both supervised (e.g., BTMs) and unsupervised (e.g., PCA, MOFA, MCIA) approaches. The diversity in methodology reflects the contestants’ attempts to capture the complex and multi-dimensional nature of the data. A critical component in the success of these models was the approach to data preprocessing. Key steps, such as normalization, handling missing values, and feature scaling, were employed by most models (19/25) to ensure the data was adequately prepared for analysis. These preprocessing techniques are known to reduce biases, standardize the data, and optimize it for model training^33^. Overall, effective data preprocessing played a crucial role in the improved performance and reliability of the predictive models.

We observed that the control models we established, which relied on the subject’s age and pre-vaccination state as task variables, performed well as baseline models for comparing more complex models submitted by contestants. Modeling post-vaccine immune responses involves significant variability due to individual differences in immune system behavior, the influence of prior exposures, and other unknown confounding factors^15,34,35^. Despite these complexities, it was essential to construct robust baseline models that captured the fundamental biological responses using minimal variables. By focusing on straightforward, readily available variables such as age and pre-vaccination state, we were able to create a reliable reference point. This allowed us to accurately assess how more complex models, incorporating immunological and demographical data, predicted post-vaccine responses. These baseline models thus played an important role in evaluating the complexity of the advanced approaches while providing a fair comparison.

The IgG-PT-D14 tasks (both titer and fold change) demonstrated the highest number of models with significant correlations, indicating that these tasks were the most successfully predicted. In contrast, the Monocyte-D1 and CCL3-D3 (tasks 2.2 and 3.2) fold change tasks had fewer models showing significant correlations, suggesting greater difficulty or variability in their prediction. Additionally, the Monocyte-D1 and CCL3-D3 (tasks 2.1 and 3.1) response prediction tasks exhibited a mix of results, with some models performing well while others demonstrated inconsistent performance, indicating moderate difficulty in predicting these tasks. These mixed results underscore the need for innovative modeling techniques to better capture the nuances of monocyte and *CCL3* responses, specifically fold-change values. Overall, the variability in prediction success across these tasks highlights the inherent challenge of modeling Tdap post-vaccination immune responses, particularly when compared to the more predictable IgG responses.

We believe this collaborative and innovative approach will create a hub for immunologists to push for novel models of immunity against Tdap boost. We expect the resultant models will also be relevant for other vaccinology studies. Contestants from the research community that are interested in participating are encouraged to contact us via and check the website (www.cmi-pb.org) for the upcoming contest information.

## Methods

### Challenge data and ground truth

The invited CMI-PB prediction challenge is outlined in **Figure 1**. A total of three multi-omics datasets were provided to contestants consisting of 117 subjects. The entire dataset was split into training and challenge datasets. The training dataset includes two independent cohorts, the 2020 cohort and the 2021 cohort, and these cohorts are discussed in detail in two recent publications: da Silva Antunes et al. ^11^ and Shinde et al.^10^, respectively. The challenge or ground truth evaluation dataset consists of 21 subjects, and we conducted experimental assays similar to those performed on the training datasets, as described in the following:

#### Experimental model and subject details

The characteristics of all 21 subjects are summarized in **Table SX3**, with human volunteers who had received either the aP or wP vaccination during childhood being recruited for the study. All participants provided written informed consent before donation and were eligible for Tdap (aP) booster vaccination. Longitudinal blood samples were collected pre-booster vaccination (day -30, -14, 0) and post-booster vaccination after 1, 3, 7, and 14 days. This study was performed with approvals from the IRB at the La Jolla Institute for Immunology, and written informed consent was obtained from all participants before enrollment.

#### Experimental data generation

Each multi-omics dataset consists of metadata about subjects and experimental data generated using four assays: plasma antibody measurements, PBMC cell frequencies, plasma cytokine concentrations, and RNA sequencing. We run experiments on three pre-booster (day -30, -14, 0) timepoints and four post-vaccine responses (day 1, 3, 7, and 14) time points.

1. **Plasma antibody measurements.** An indirect serological assay was employed using xMAP Microspheres (Luminex Corporation) to measure TdaP antigen-specific antibody responses in human plasma. Pertussis antigens (PT, PRN, Fim2/3, FHA), Tetanus Toxoid (TT), Diphtheria Toxoid (DT) , and Ovalbumin (negative control) were coupled to uniquely coded beads (xMAP MagPlex Microspheres). A detailed description is provided by da Silva Antunes et al.^11^.
2. **PBMC cell frequencies.** Twenty-one different PBMC cell subsets were identified using manual gating using FlowJo (BD, version 10.7.0). The detailed description is provided here^11^.
3. **Plasma cytokine concentrations.** Plasma samples were randomly distributed on 96 well plates for quantification of different plasma cytokines by Olink proteomics assay. The detailed description is provided here^11^.
4. **RNA sequencing.** Library preparation was performed using the TruSeq Stranded mRNA Library Prep Kit (Illumina). Libraries were sequenced on a HiSeq3000 (Illumina) system. The paired-end reads that passed Illumina filters were further filtered for reads aligning to tRNA, rRNA, adapter sequences, and spike-in controls. The remaining reads were aligned to the GRCh38 reference genome and Gencode v27 annotations using STAR (v2.6.1)^36^. DUST scores were calculated with PRINSEQ Lite (v0.20.3)^37^, and low-complexity reads (DUST >4) were removed from the BAM files. The alignment results were parsed via the SAMtools to generate SAM files^38^. Read counts to each genomic feature were obtained with the featureCounts (v1.6.5 using the default options along with a minimum quality cut-off (Phred >10))^39^.

Contestants were supplied with the baseline immunoprofiling data for all challenge dataset subjects. The post-vaccine response data, which contain the ground truth, were hidden from the contestants.

#### Data processing

In addition to the original raw data generated by immunoprofiling, we performed data pre-processing as described in **Figure 1**. In addition to the original raw data generated by immunoprofiling, we performed data pre-processing as described in **Figure 1**. First, we identified common features between the training and challenge datasets and excluded features with a coefficient of variance less than 0.3. Second, we performed baseline normalization on cell frequency, antibody titer, and cytokine concentration data. Specifically, we calculated the baseline median as a normalization factor per analyte and divided all values by this factor. Third, we ran CombatSeq with default parameters to correct batch effects^12^. To maintain consistency, we performed baseline normalization on cell frequency, antibody titer, and cytokine concentration data in the test dataset but did not apply any normalization to the gene expression data.

### Formulating the prediction tasks

Contestants were challenged to predict a ranked list of the highest response (to be ranked first) to the lowest response (to be ranked last) subjects for each prediction task provided. We formulated six prediction tasks in order to quantitatively compare different approaches to model immune responses to Tdap booster vaccination. We selected biological readouts known to be altered by booster vaccination under the premise that these readouts would likely capture meaningful heterogeneity across study subjects based on our previous work^11^. We formulated six prediction tasks: three required contestants to predict specific biological readouts on particular days following the vaccine response, and the other three required contestants to predict the fold change between specific biological readouts on particular days following the vaccine response and the pre-vaccination state.

In task 1.1, contestants had to predict plasma IgG levels against the pertussis toxin (PT) on day 14 post-booster vaccination. For task 1.2, contestants were required to predict the fold change of the plasma IgG levels against the pertussis toxin (PT) between day 14 post-booster vaccination and baseline. Tasks 2.1 and 2.2 required contestants to predict the overall frequency of monocytes among PBMCs on day 1 post-booster vaccination and the corresponding fold change, respectively. Similarly, tasks 3.1 and 3.2 required contestants to predict CCL3 gene expression on day 3 post-booster vaccination and the corresponding fold change values compared to baseline.

### Prediction challenge evaluation

After receiving contestants’ predicted ranked list for each task, we curated the rank file. If we found NA values in the ranked list, we imputed them with the median rank for that list. Evaluations were performed in two steps.

First, we chose the Spearman rank correlation coefficient as evaluation metric to compare the predicted ranked list (*p*) for each task, *t*, and n subjects (n=21 for the set of challenge dataset subjects), *R_p,t_ = (r_p,1_, r_p,2_, …, r_p,n_)* against ground truth (*g*) ranked list *R_g,t_ = (r_g,1_, r_g,2_, …, r_g,n_)*. The Spearman rank correlation coefficient (ρ) is given by:

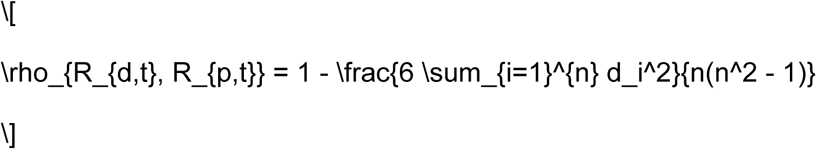

where *d_i_ = R_g,i_* - *R_p,i_* is the difference between the ranks of each pair. In this way, each task submitted by constant was evaluated.

Second, we devised a point system to rank all submissions and identify the overall winner of the challenge. Specifically, we awarded 3 points if a submission was top-ranked in a particular task and 1 point if the contestant attempted the task.

## Data availability

Training and challenge data of this prediction challenge can be found at: https://www.cmi-pb.org/data

## Quantification and statistical analysis

Statistical analyses are detailed for each specific technique in the specific Methods section or in the figure legends, where each specific comparison is presented. Statistical tests were performed using R (version 4.1, www.r-project.org/) of the Spearman correlation coefficient. Details pertaining to significance are also noted in Figure 2 legends, and p < 0.05 is defined as statistical significance.

## Limitations of the study

Our challenge dataset cohort comprised multi-omics data for 21 subjects, a size smaller than the training data cohorts of 96 subjects, maintaining just an 80:20 training-to-challenge dataset ratio. The smaller size of the challenge cohort may result in reduced precision and heightened sampling variation in Spearman rank calculations, which were used as an evaluation matrix, potentially impacting the reliability and generalizability of correlation results. However, models developed by contestants exhibited strong performance, surpassing control models in four tasks. To address this limitation, we intend to enhance the size of our challenge cohort in the upcoming public contest.

## Acknowledgments

We are grateful to the La Jolla Institute for Immunology Flow Cytometry and Bioinformatics core facilities for their services. The authors would like to thank all donors who participated in the study and the clinical studies group staff - particularly Gina Levi - for all the invaluable help. Research reported in this publication was supported by the National Institute of Allergy and Infectious Diseases of NIH under award nos. U01AI150753, U01AI141995, and U19AI142742.

## Author Contributions

All contestants were included as co-authors on this manuscript.

**Supplementary Information:** Enclosed with this document

**Supplementary Note 1:** A detailed description of CMI-PB invited prediction challenge methods.

