## Supplementary Information for "Putting computational models of immunity to the test - an invited challenge to predict *B. pertussis* vaccination outcomes"

**Summary of feedback received after the conclusion of the challenge:**

After completing the invited participant challenge, we asked participants to provide their feedback on the challenge. We sent participants a poll with questions regarding the resources, the difficulty level of the prediction tasks, future participation, and suggestions to help. The results of the feedback poll indicated that the most useful resource to participants was direct communication with the CMI-PB Consortium members (ie. emailing) and the least useful resource was the three Zoom Informational Sessions, which were held to demonstrate the submission process and the website live, as well as to encourage participants to ask questions to the CMI-PB Team in real-time.

When asked about the difficulty of the prediction tasks, with 1 being very difficult and 6 being very simple, the average response was 3.22. This showed us that the prediction tasks were evenly balanced and they were not too simple, yet not too challenging. When asked if they would participate in a similar challenge, the average response was 5.55, with 1 being very unlikely and 6 being very likely. The participants were also asked about their overall satisfaction with the challenge, and we received an average score of 5.22, with 1 being very dissatisfied and 6 being very satisfied. Overall, the feedback poll indicated that participants were satisfied with the overall experience of the prediction challenge and are looking forward to the 3rd (public) challenge.

This invited CMI-PB challenge has been designed to address some of the shortcomings identified during the first challenge. Based on the second challenge, we expect to make additional adjustments to help ensure success in the initial public challenge. This iterative process aims to provide contestants with a rich user experience, allowing for smoother data access and a much less tedious prediction submission process.

### Supplementary Tables

**Table S1**. Antibody information.

| Target | Conjugate | Host | Target | Clone | Catalog | Vendor | Dilution |
|---|---|---|---|---|---|---|---|
| CD45 | BUV395 | Mouse | Human | HI30 | 563792 | BD | 1/500 |
| CD8 | BUV496 | Mouse | Human | RPA-T8 | 612942 | BD | 1/500 |
| CD20 | BUV563 | Mouse | Human | 2H7 | 748456 | BD | 1/200 |
| CD16 | BV510 | Mouse | Human | 3G8 | 612786 | BD | 1/100 |
| CD3 | BUV805 | Mouse | Human | UCHT1 | 612895 | BD | 1/200 |
| CD14 | BV480 | Mouse | Human | M5E2 | 746304 | BD | 1/100 |
| CD45RA | BV570 | Mouse | Human | HI100 | 304132 | Biolegend | 1/200 |
| CD19 | BV605 | Mouse | Human | HIB19 | 302244 | Biolegend | 1/200 |
| IgD | PE-CF594 | Mouse | Human | IA6-2 | 747484 | BD | 1/200 |
| CD11c | BV785 | Mouse | Human | 3.9 | 301644 | Biolegend | 1/50 |
| CCR7 | FITC | Mouse | Human | G043H7 | 353216 | Biolegend | 1/66 |
| CD123 | PE-Cy7 | Mouse | Human | 6H6 | 306016 | Biolegend | 1/100 |
| CD38 | PerCP-Cy5.5 | Mouse | Human | HIT2 | 562288 | BD | 1/50 |
| HLA-DR | AF700 | Mouse | Human | L243 | 307616 | Biolegend | 1/50 |
| CD56 | APC | Mouse | Human | 5.1H11 | 362504 | Biolegend | 1/100 |
| CD4 | APC-eF780 | Mouse | Human | RPA-T4 | 47-0049-42 | LIFE TECH | 1/200, 1/50 |
| CD71 | PE-Cy5 | Mouse | Human | M-A712 | 551143 | BD | 1/50 |
| CD66b | BV421 | Mouse | Human | G10F5 | 562940 | BD | 1/100 |
| CD1c | BV650 | Mouse | Human | 1.161 | 331542 | Biolegend | 1/200 |
| CD141 | PE | Mouse | Human | M80 | 47-0049-42 | LIFE TECH | 1/200 |

**Table S2**. The characteristics of all 21 subjects in the challenge dataset.

| Subject ID | Age | Biological Sex at Birth | Vaccine Priming Status |
|---|---|---|---|
| 97 | 35 | Male | wP |
| 98 | 28 | Female | wP |
| 99 | 22 | Female | aP |
| 100 | 20 | Female | aP |
| 101 | 18 | Male | aP |
| 102 | 18 | Male | aP |
| 103 | 27 | Female | wP |
| 104 | 32 | Female | wP |
| 105 | 27 | Female | wP |
| 106 | 25 | Female | aP |
| 107 | 23 | Female | aP |
| 108 | 26 | Female | wP |
| 109 | 32 | Female | wP |
| 110 | 24 | Female | aP |
| 111 | 25 | Male | wP |
| 112 | 25 | Male | aP |
| 114 | 31 | Male | wP |
| 115 | 19 | Female | aP |
| 116 | 21 | Male | aP |
| 117 | 27 | Female | aP |
| 118 | 23 | Male | aP |

### Supplementary Figures

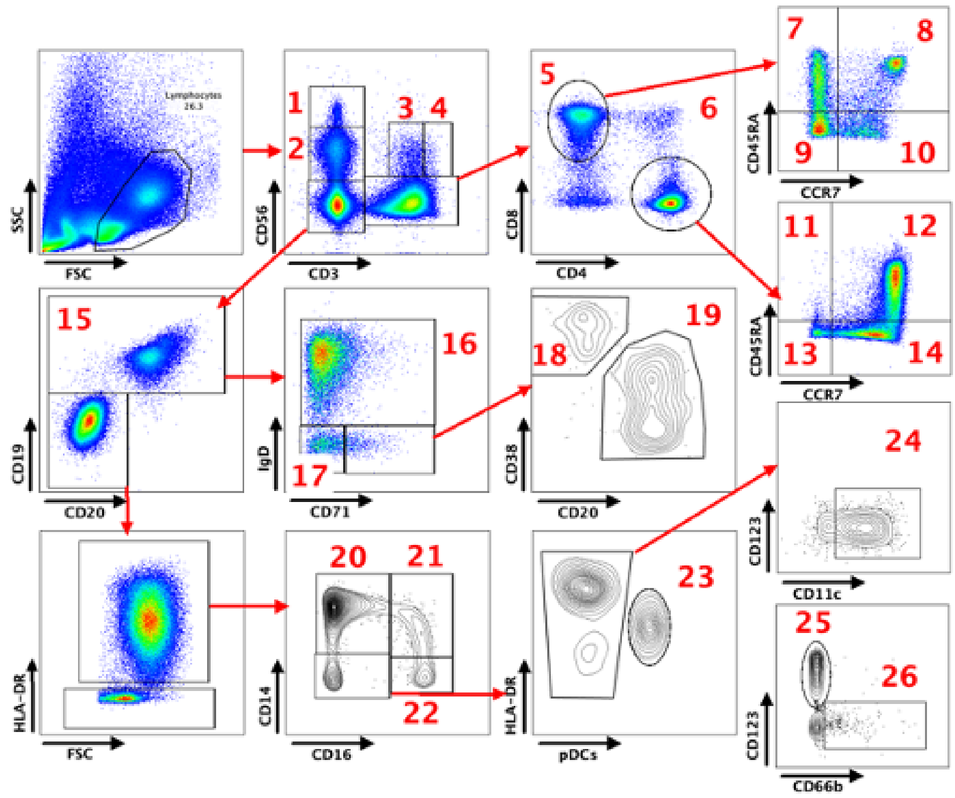


| # | Cell population |
|---|---|
| 1 | CD56$^{high}$ NK cells |
| 2 | CD56$^{+}$ NK cells |
| 3 | CD56$^{+}$ T cells |
| 4 | CD56$^{+}$ CD3$^{high}$ T cells |
| 5 | CD8 T cells |
| 6 | CD4 T cells |
| 7 | CD8 TEMRA |
| 8 | CD8 Naïve |
| 9 | CD8 TEM |
| 10 | CD8 TCM |
| 11 | CD4 TEMRA |
| 12 | CD4 Naïve |
| 13 | CD4 TEM |
| 14 | CD4 TCM |
| 15 | B cells |
| 16 | Naïve B cells |
| 17 | Memory B cells |
| 18 | Plasmablasts |
| 19 | Activated B cells |
| 20 | Classical Monocytes |
| 21 | Intermediate Monocytes |
| 22 | Non-classical Monocytes |
| 23 | pDCs |
| 24 | Conventional DCs |
| 25 | Basophils |
| 26 | Activated granulocytes |

**Figure S1**. Gating strategy PBMC cell frequencies (FACS).

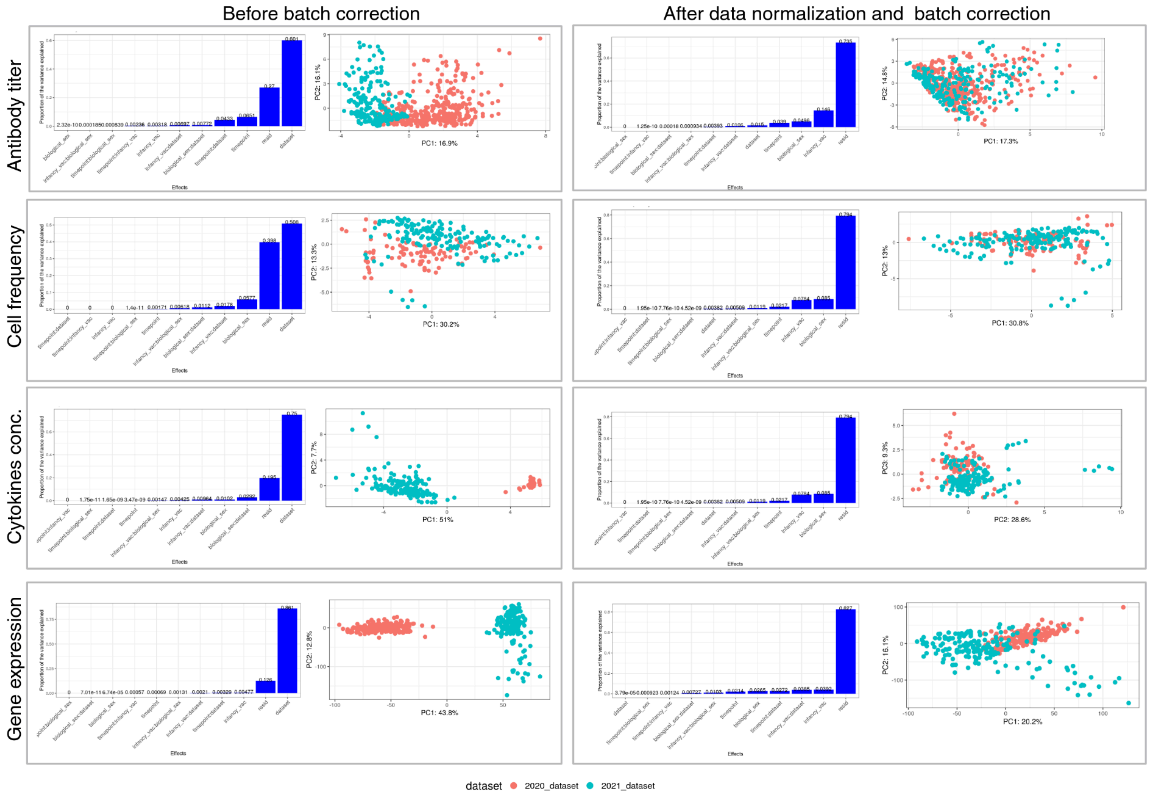


**Figure S2: Plot of assay data before and after normalization and batch effect correction**. For each assay, the plots on the left represent data before batch correction, while the plots on the right represent data after normalization and batch correction.
